# DNA sequence properties that predict susceptibility to epiallelic switching

**DOI:** 10.1101/057794

**Authors:** Marco Catoni, Jayne Griffiths, Claude Becker, Nicolae Radu Zabet, Carlos Bayon, Mélanie Dapp, Michal Lieberman-Lazarovich, Detlef Weigel, Jerzy Paszkowski

## Abstract

Transgenerationally heritable epialleles are defined by the stable propagation of alternative transcriptional states through mitotic and meiotic cell cycles. Given that the propagation of DNA methylation at CpG sites, mediated in Arabidopsis by MET1, plays a central role in epigenetic inheritance, we examined genome-wide DNA methylation in partial and complete loss-of-function *met1* mutants. We interpreted the data in relation to transgenerational epiallelic stability and provide evidence that DNA sequence features such as density of CpGs and genomic repetitiveness can be used to predict susceptibility to epiallelic switching. The importance of these rules was confirmed by analyses of common epialleles in natural Arabidopsis accessions and verified in rice.

## Introduction

Transgenerational epigenetic inheritance have been well documented in plants, however the primary competence that leads to the formation of alternative epialleles at only certain loci is not well understood. It has been reported that maintenance of ^m^CpG (methylated CpG) patterns by MET1 and the chromatin remodeler DDM1 is central for transgenerational epigenetic inheritance (Reinders et al. 2009; Teixeira et al. 2009). Importantly, inactivation of MET1 or DDM1 results in the loss of ^m^CpGs, which is not easily corrected after re-introduction of MET1 and DDM1, although remethylation occurs at certain loci (Reinders et al. 2009; Teixeira et al. 2009). Thus, chromosomal targets of CpG methylation were divided into two broad categories: (a) those that can form two distinct epigenetic states (epialleles) that are maintained over generations in the presence of all epigenetic activities of the wild type, and (b) those that revert to only one dominant epigenetic state (reversible) and are thus not able to form heritable epialleles (Reinders et al. 2009; Teixeira et al. 2009). Although epiallelic reversion was associated with RNA directed DNA methylation (RdDM)(Teixeira et al. 2009), the primary determinants underlying differences between the chromosomal targets of epigenetic regulation in susceptibility to epiallelic switching remained unknown. Here we compare the whole-genome distribution of DNA methylation in Arabidopsis plants carrying either a weak or strong allele of *MET1*. Although the partial loss-of-function allele *met1-1* reduces CpG methylation levels to approximately 25% of wild type (Kankel et al. 2003), this causes only minor developmental defects. In contrast, the null allele *met1-3* causes an almost complete loss of ^m^CpGs and is semi-lethal (Mathieu et al. 2007). The methylation remaining in *metl-1* thus identifies a particularly important subset of ^m^CpG sites, offering a unique opportunity to understand the function of ^m^CpG methylation. We find that loci forming stable epialleles are similarly affected in the two mutants, while epigenetically reversible loci are affected differently in *metl-1* and *metl-3*. These observations allow inferences about the molecular mechanisms of epigenetic transgenerational inheritance at these distinct classes of loci and the formulation of genetic and epigenetic rules predicting the capacity of chromosomal targets to form stable epialleles.

## Results and Discussion

While the null allele *met1-3* causes an almost complete loss of ^m^CpGs (Lister et al. 2008; Tariq et al. 2003) and is semi-lethal (Mathieu et al. 2007), the partial loss-of-function allele *met1-1*, which reduces CpG methylation levels to approximately 25% of wild type (Kankel et al. 2003), causes only minor developmental defects. This suggests that the 25% ^m^CpGs remaining in *met1-1* (Figure S1) are important for a large fraction of epigenetic information. Therefore, we compared whole-genome transcriptomes and methylomes between wild-type Col-0 plants, *met1-1* and *met1-3*, and also the *F2* progeny of a hybrid derived from a cross between *met1-3* and wild type. The *F2* plants were genotyped and only individuals with homozygous wild-type alleles of *MET1* were analysed. These *MET1*^+^ *F_2_* (hereafter *MET1*^+^) siblings had inherited half of their genomes from a *met1-3* grandparent, except for the region on chromosome 5 around *MET1.* For simplicity, we excluded chromosome 5 from subsequent analyses. The *MET1*^+^ segregants had on average 57% of wild-type ^m^CpGs that indicated remethylation of the sequences inherited from the met1 grandparent (Figure S1C, Table S1).

Next, we screened for differentially methylated regions (DMRs) in pairwise comparisons with *met1-3*, *metl-1* and *MET1+* plants using wild type as the common denominator. DMRs in *met1-3* included almost all DMRs of both *met1-1* and *MET1^+^* plants (Figures 1A and S2). Interestingly, DMRs of *met1-1*and *MET1^+^* segregants overlapped in more than 85′000 commonly methylated cytosines, representing 57% and 48% of *met1-1* and *MET1^+^* DMRs, respectively (Figure 1B). There was a significant correlation (Spearman R^2^=0.48, Pearson R^2^=0.63) of methylation distribution between *metl-l* and *METl^+^* (Figure 1C), which was further confirmed by hierarchical clustering of ^m^CpGs in 200-bp non-overlapping genomic windows (tiles) (Figure 1D). These results suggested that certain methylation patterns in *met1-1* and *MET1^+^* are associated with particular loci, possibly due to intrinsic characteristics of DNA sequences of the affected loci themselves. This hypothesis was tested in subsequent experiments.

**Figure 1.**
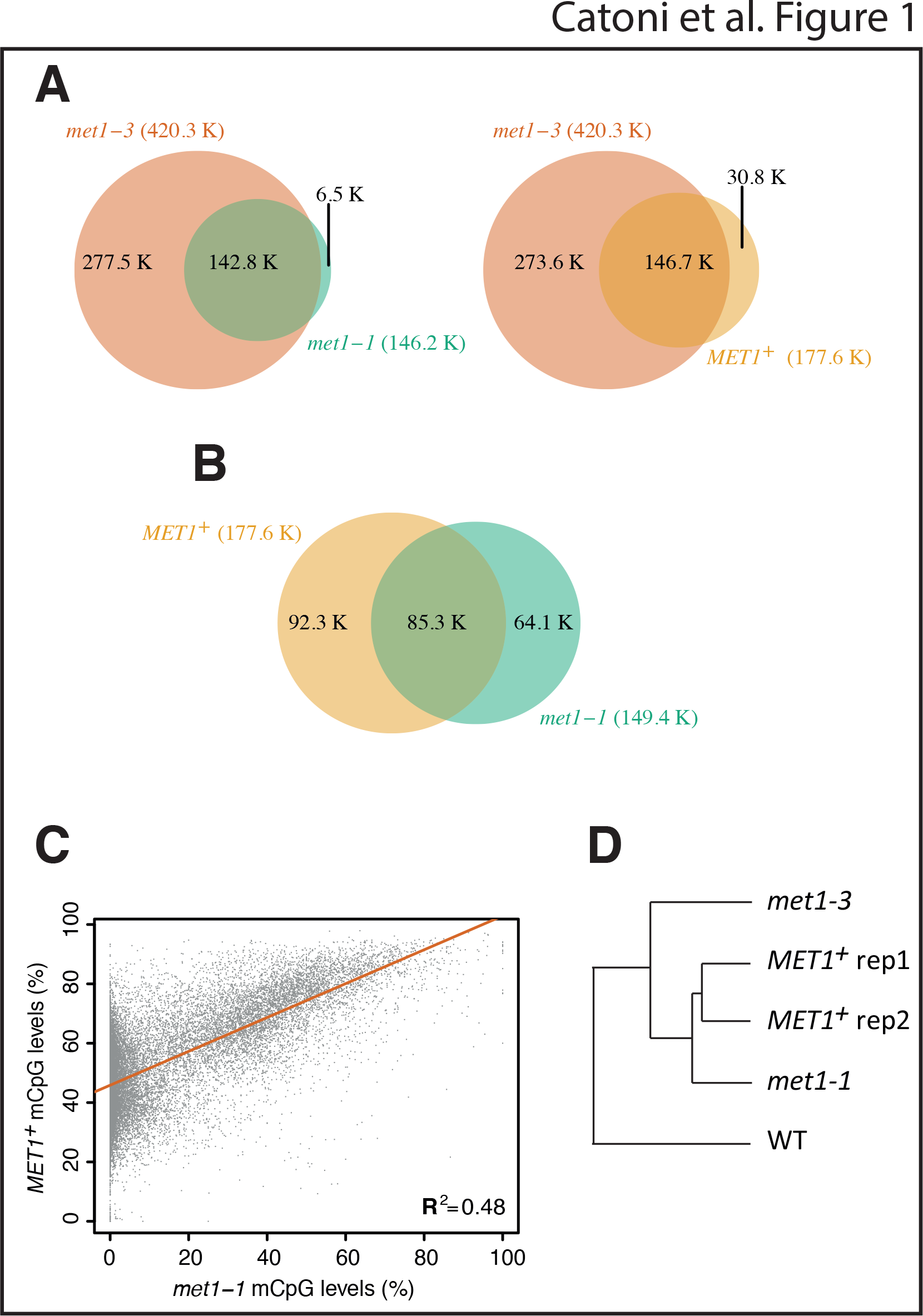
Patterns of CpG methylation in *met1-1* and *MET1^+^* plants.

(A) Venn diagrams of genome-wide hypomethylated CpG DMRs (chromosomes 1 to 4) of *met1-3* compared to control Col-0 plants and later compared to DMRs of *met1-1* and *MET1^+^*. For *metl* and *MET1^+^* plants, DMRs were considered with a relative difference to wild type of ≥80% and ≥40%, respectively. Overlapping areas represent number of cytosines in shared portions of DMRs.

(B) Venn diagram of genome-wide hypomethylated CpG DMRs (chromosomes 1 to 4) of **metl-1** (≥80% difference) and combined (union of DMRs) *MET1^+^* plant pools (≥40% difference). Permutation test with 1000 iterations using regioneR package(Gel et al. 2016) confirmed a statistically significant overlap (p-value=0.001).

(C) Scatter plot of CpG DMRs (chromosomes 1 to 4) of *metl-1* and union of DMRs *MET1^+^* plants (DMR criteria as in A and B). Linear regression - red line. The correlation coefficient is indicated in the bottom right corner of the plot.

(D) Hierarchical clustering of CpG methylation in genomic tiles of wild type (wt), *metl-1*, *metl-3* and *MET1^+^* plants (chromosomes 1 to 4). The methylation of *MET1^+^* was adjusted by subtracting 50% of the methylation derived from wild type. Clustering based on randomly selected genomic tiles (10%) (*n*=13,668) representing ≥50% CpG methylated in wild type were used for analyses.

It has been proposed that targets of DNA methylation in plants are of two distinct types (Saze and Kakutani 2011): (1) gene body methylation (hereafter referred to as Gene Body Like or GEL), present mostly in coding regions of expressed genes and consisting exclusively of ^m^CpGs, and (2) transposable element methylation (hereafter referred to as Transposable Element Like or TEL), found predominantly at transposons and chromosomal repeats and affecting CpGs but also non-CpGs. Here, GELs were defined as gene models in which averaged methylation of cytosines in CpG context was above 5%, but less than 5% in non-CpG contexts and TELs as loci with methylation of both ^m^CpGs and non-CpGs over 5% (Table S2). This produced totals of 11,746 GELs and 4,743 TELs. As expected, 98% of GELs were annotated as genes, while 73% of TELs were annotated as transposable elements (Figure S3) and 22% as genes (Figure S3), indicating that transposon-like methylation is also associated with a subset of protein coding genes. We also annotated this way 200-bp genomic tiles containing at least 5% of averaged CpG methylation. We then examined GEL and TEL methylation in *met1-3*, *met1-1* and *MET1^+^*. In *met1-3*, the two types of methylation were equally erased (Figure S4). In contrast, although in *met1-1* methylation at GELs was uniformly lost, methylation losses were not uniform across TELs, with many predominantly losing and others predominantly maintaining DNA methylation (Figures 2A and S5). Interestingly, *MET1^+^* segregants displayed a methylation pattern very similar to *met1-1*. Analyses of GELs in the *MET1^+^* segregants revealed an average methylation of 50%, suggesting ubiquitous maintenance of the mid-parental methylation levels (Figures 2A, S4 and S5) and TELs in the *MET1^+^* segregants revealed an average methylation of 70% relative to wild type, implying substantial remethylation above the mid-parental level (Figures 2A, S4 and S5). Notably, the patterns of methylation at particular TELs in *met1-1* and *MET1^+^* were overlapping (Figures 2A and S5), indicating that TELs have similar remethylation capacities in the two genotypes or, in the case of *met1-1*, similar resistance to methylation loss. The tiling arrays transcriptome mapping of *metl-1*, *metl-3* and wild type revealed that residual methylation in **metl-1** was also reflected by transcriptional silencing of these TELs (Figure S6).

**Figure 2.**
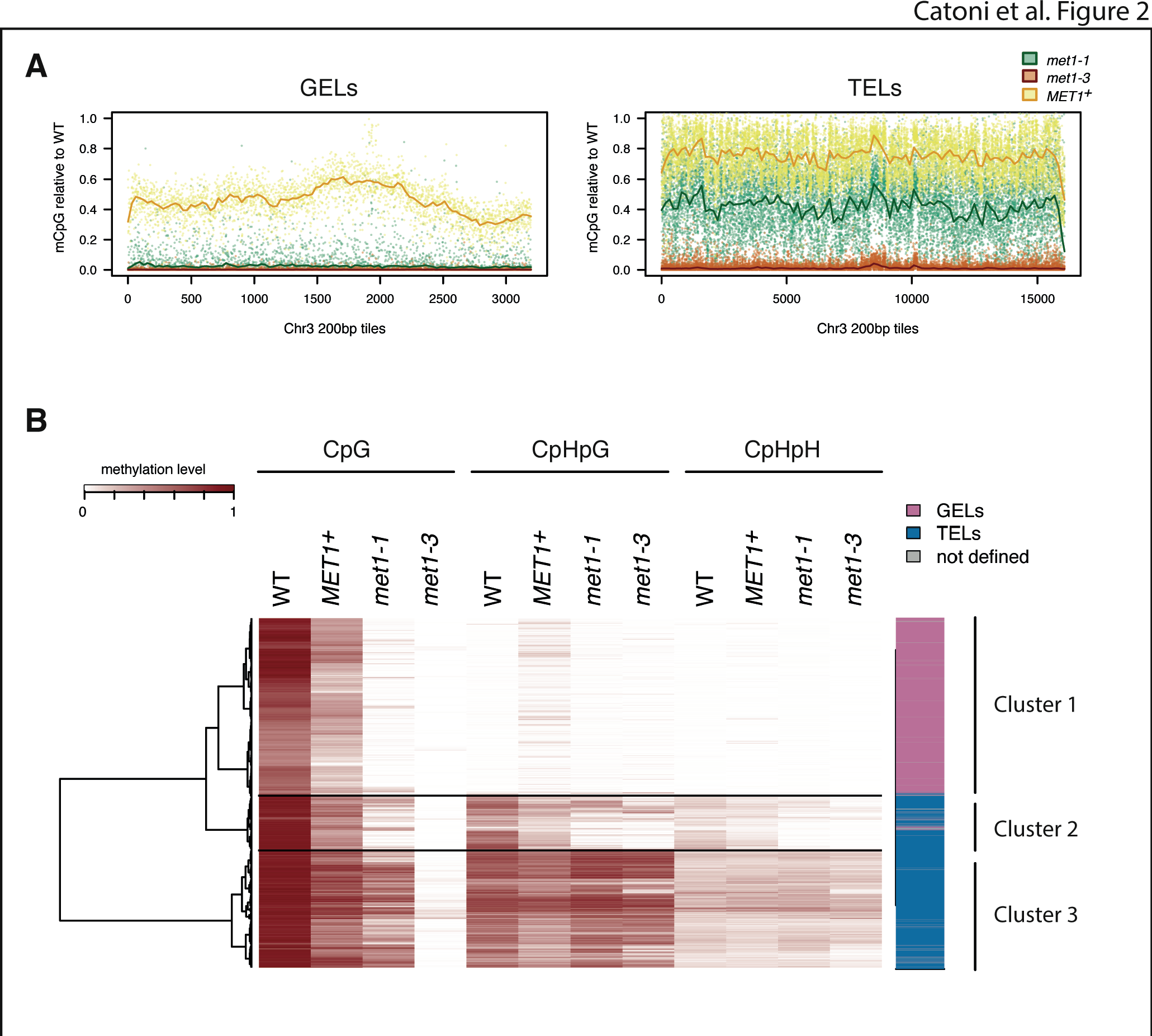
Distribution of CpG methylation at GELs and TELs in wild type, *metl-1*, *metl-3* and *MET1^+>^*.

(A) Distribution of CpG methylation levels at 200-bp tiles (≥80% CpG methylated in wild type) along chromosome 3 at GELs or TELs (Table S2). Lines were generated using lowest smooth R with a span of 1/50. Legend to colours assigned to particular genotypes is above the right graph. *MET1^+^* showed is mean value of two replicates. Corresponding graphs for chromosomes, 2, and 4 are in Figure S5.

(B) Heatmap R of DNA methylation levels at 200-bp tiles (10% randomly selected of chromosome 1, CpG methylated in wild type ≥40%, *n*=11,094). Sequence context of methylation, genotypes and methylation levels are indicated above the figure. Tiles were ordered according to hierarchical clustering (hclust, R software environment) and the main clusters are delineated by horizontal lines and explained at the right of the figure and in the main text. The Euclidean distance dendrogram is presented left of the figure. *MET1^+^* sample is derived from the second replicate.

Using 200-bp tiles, we further classified CpG-methylated regions according to their association with methylation of non-CpGs and performed hierarchical clustering that defined three main clusters (Figure 2B). The cluster 1 contained almost exclusively GELs, displaying the greatest methylation deficiency in *MET1^+^* segregants relative to other clusters, while cluster 2 and cluster 3 identified TELs with different degree of methylation in *MET1^+^* and **metl-1** (Figures 2B and S7). Further comparison of the global CpG methylation levels of GELs and TELs in *met1-3*, *met1-1* and *MET1^+^* in relation to wild type indicated that in **met1-3**, GELs and TELs both lacked ^m^CpGs, while in *met1-1* and *MET1^+^*TELs had retained or regained significant levels of CpG methylation (Figure 2B and S4). These methylome and clustering analyses suggested that TELs are heterogeneous, consisting of at least two classes (Figure 2B).

To define genetic and epigenetic features of TELs that result in differential methylation losses in *met1-1*, correlated with methylation levels observed in *MET1^+^*, we sought DNA sequence properties of TELs that coincide with persistence or loss of CpG methylation in *met1-1*. We discovered that increase in the number of CpGs is associated with the propensity of TELs to lose CpG methylation (Figure 3A), suggesting a link between DNA sequence properties at methylated TELs and the formation of stably demethylated epialleles. We also found that presence of tandem repeats longer than 100 bp and increasing levels of their repetitiveness jointly correlate with maintenance of methylation in *met1-1* or thus remethylation in *MET1^+^* (Figure 3B). Therefore, intrinsic features of TEL DNA sequence are associated with the distribution of their methylation in *met1-1* and also with the formation of transgenerationally stable methylation patterns (epialleles) in *MET1^+^* segregants.

**Figure 3.**
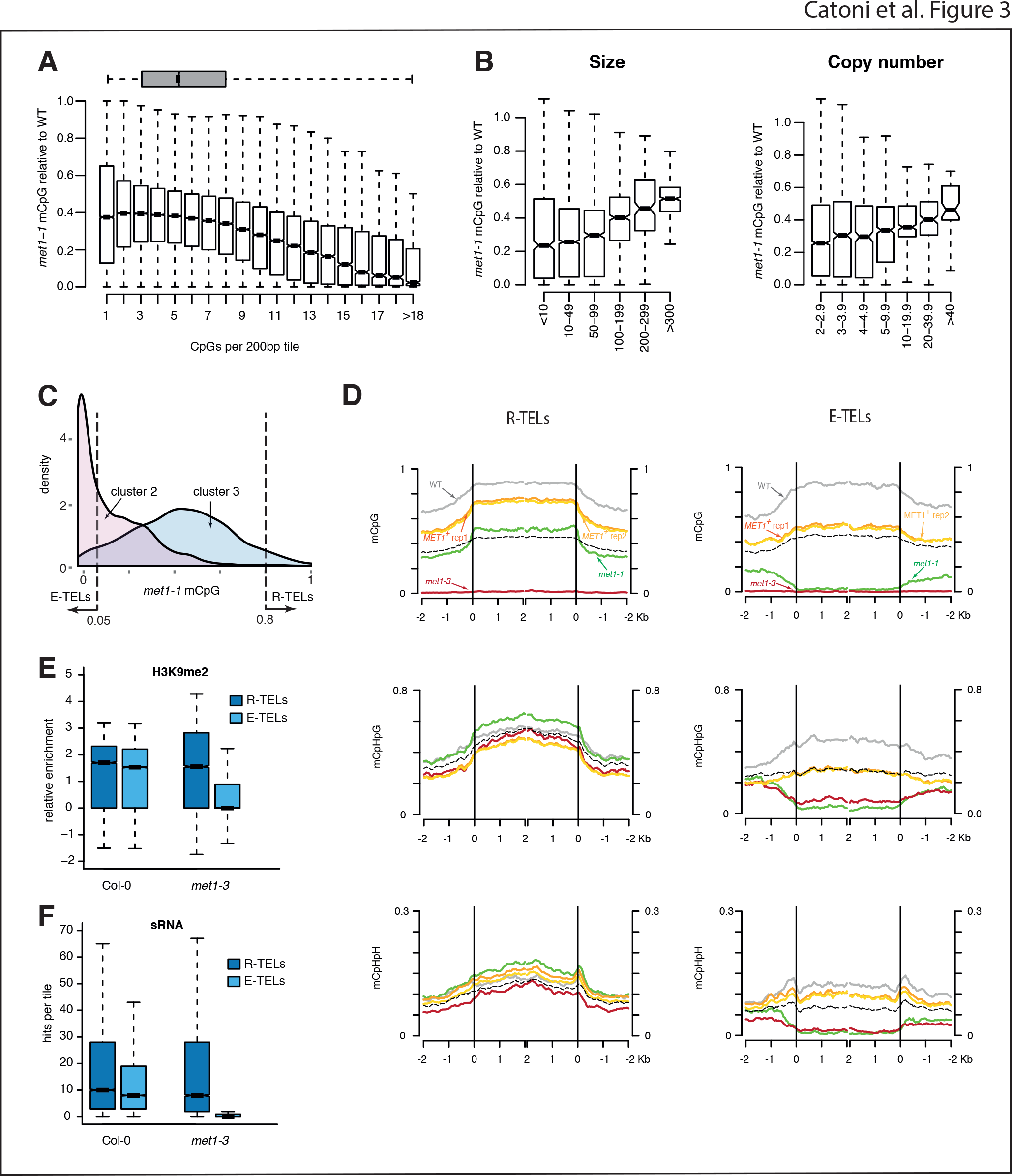
Genetic and epigenetic properties of TELs.

(A) Box plot of CpG methylation levels in *met1-1* relative to wild type (wt) in 200-bp tiles assigned to TELs (Table S2) and sorted according to the number of CpGs. The genome-wide tile distribution is illustrated by a grey box plot at the top of the chart.

(B) Box plot of CpG methylation levels in *met1-1* relative to wild type (wt) in tandem repeats sorted according to the size of the repeat units (left) or their copy number (right).

(C) Density plot of CpG methylation level distribution in *met1-1* relative to wild type for 200-bp tiles assigned to sub-cluster 2 and cluster 3 as defined in Figure 1B. Vertical dashed lines mark criteria for E-TELs and R-TELs selection which incorporate mostly tiles from cluster 2 and cluster 3, respectively.

(D) Kernel density plots of DNA methylation levels in wild type, *met1-1*, *met1-3* and *MET1^+^* plants at R-TELs and E-TELs. Identities of R-TEL and E-TEL (defined in C) are according to TAIR10 annotation and aligned at the 5′ end or the 3′ ends (vertical lines). Average methylation levels for each 200-bp window are plotted. R-TELs and E-TELs at chromosome 5 were excluded from the analyses. Black dashed line displays WT and *met1-3* mid-parental methylation.

(E) Box plot of H3K9me2 levels at R-TELs and E-TELs in wild type and *met1-3* (Deleris et al. 2012). Analyses were performed with 200-bp windows.

(F) Box plot of small RNA levels at R-TELs and E-TELs in wild type and *met1-3* (Lister et al. 2008). Analyses were performed with 200-bp windows.

To further characterize the epiallelic behaviour of TELs, we rank ordered TELs of cluster 2 and 3 (Figure 2B) according to their levels of methylation in *met1-1* and selected two contrasting subsets of TELs for further study (Figures 3C and S8). The first subset retained less in *met1-1* than 5% of wild-type CpG methylation. As this subset resembled GELs in the capacity to form stable epialleles, we refer to them as Epiallelic-TELs or E-TELs. More than 80% of CpG methylation was retained in *met1-1*in the second subset, correlating with regain of CpG methylation in *MET1^+^* and, thus, rapid reversal of the levels towards wild type. This subset of TELs was named Reversible-TELs or R-TELs. It is important to bear in mind that GELs, E-TELs and R-TELs are equally depleted of CpG methylation in *metl-3* (Figure S4).

To directly compare methylation levels at all cytosines of E-TELs and R-TELs in wild-type, *met1-3*, *met1-1* and *MET1^+^* plants, we aligned their annotated sequences to construct plots for CpG, CpHpG and CpHpH methylation (Figure 3D). E-TELs displayed a complete loss of CpG methylation in both met1 mutants and showed midparent levels of 50% of wild type in *MET1^+^* segregants, suggesting a lack of or minimal remethylation activity at these sequences by MET1, which is present already in *met1-3*/*MET1^+^* F1 hybrids and in *MET1^+^* segregants. In contrast, although R-TELs completely lost CpG methylation in *met1-3*, they reached approximately 80% of wild-type ^m^CpGs in *MET1^+^* segregants (Figure 3D), supporting their active remethylation in the presence of MET1. Interestingly, although E-TELs and R-TELs do not show a relevant difference in their non-CpG methylation levels in wild-type plants (Figure S9), both CpHpG and CpHpH methylation were significantly reduced in both *met1* alleles at E-TELs but not at R-TELs (Figure 3D). Moreover, in *MET1^+^* plants, only E-TELs displayed a significant reduction in CpHpG methylation compared to the wild type (Figure 3D). Thus, non-CG methylation seems to persist at R-TELs in both *met1* mutant alleles but is depleted at E-TELs.

Methylation at CpHpGs is a part of the self-reinforcing regulatory loop with histone 3 dimethylation in lysine 9 (H3K9me2), and methylation at CpHpHs is maintained primarily by the RdDM pathway directed by small RNAs (Law and Jacobsen 2010). Using published genomic data for H3K9me2 and sRNA distribution data available for wild type and *met1-3* (Deleris et al. 2012; Lister et al. 2008), we surveyed their distributions over R-TELs and E-TELs (Figure 3E, 3F). In wild-type plants, the levels of both H3K9me2 and the different classes of small RNAs were similar at R-TELs and E-TELs; however, in the *met1-3* mutant only R-TELs retained near wild-type levels of both H3K9me2 and sRNAs, while E-TELs lose both (Figures 3E, F and S10). These results are consistent with previously observed association of sRNAs and non-CpG methylation with transgenerationally “remethylable” loci (Teixeira et al. 2009).

Next, we examined whether E-TELs and/or R-TELs form larger epigenetically co-regulated domains or their local DNA sequences determine susceptibility to epigenetic switching. We found multiple examples of neighbourhoods of R-TELs and E-TELs (Figure S11), consistent with the hypothesis that local features of DNA sequences can be used for the prediction of epigenetic reversibility (R-TELs) or ability to form stable epialleles (E-TELs) (Figure 3A and 3B). Certain transposon superfamilies are overrepresented among E-TELs or R-TELs (Figure S12), the most striking of which are Helitrons, which have the lowest CpG content and are present exclusively in R-TELs (Figure S12). Moreover, although most TE superfamilies were represented in E-TELs and R-TELs, certain TE families were enriched differentially among E-TELs and R-TELs (Tables S3, S4 and Figure S13A), indicating that in addition to the sequence characteristics of a particular family, further sequence features, as delineated here, may explain the epigenetic properties of TEs. Certain transposon families could be clearly separated, consistent with E-TEL or R-TEL characteristics (Figure S13B), thus further reinforcing the hypothesis that DNA sequence composition, in combination with repetitiveness, can be used in defining likelihood of epiallelic properties of loci. Interestingly, one TE family (ATENSPM5) was represented in both E-TELs and R-TELs (Figure S13A). This unusual feature of ATENSPM5 appeared to exhibit a surprising duality of epigenetic regulation within this transposons by which one open reading frame (ORF) behaves like an E-TEL and the other as an R-TEL (Figure S14), again supporting the importance of very local DNA sequence determinants in differential epigenetic regulation.

To directly determine transgenerational epigenetic properties of GELs, E-TELs and R-TELs, and especially the remethylation timing of R-TELs, we backcrossed *met1-3*, which is in the Col-0 accession, to the wild-type Landsberg *erecta*(Ler) accession. This allowed the use of DNA sequence polymorphism to discriminate between alleles of the two parents in F1 hybrids and, thus, to separately examine their methylation levels. Parental methylation levels of GELs and E-TELs were maintained in F1 plants, displaying clear epi-heterozygosity (Figure 4A). In contrast, *met1-3*-derived R-TELs underwent efficient remethylation, suggesting that de-*novo* DNA methylation occurred as soon as functional MET1 became available (Figure 4B). Therefore R-TELs are clearly different from “remethylable” loci where remethylation does not occur in the F1(Teixeira et al. 2009). In addition, we examined in reciprocal backcrosses expression at R-TELs and E-TELs. The expression of E-TELs observed in *met1-1* was maintained in F1 plants, R-TEL expression was efficiently silenced to the initial wild-type level, independent of the crossing direction (Figure 4C, D). This was true for loci annotated as transposons as well as for protein-coding genes with R-TELs or E-TELs in their promoters (Figure 4C, D).

**Figure 4.**
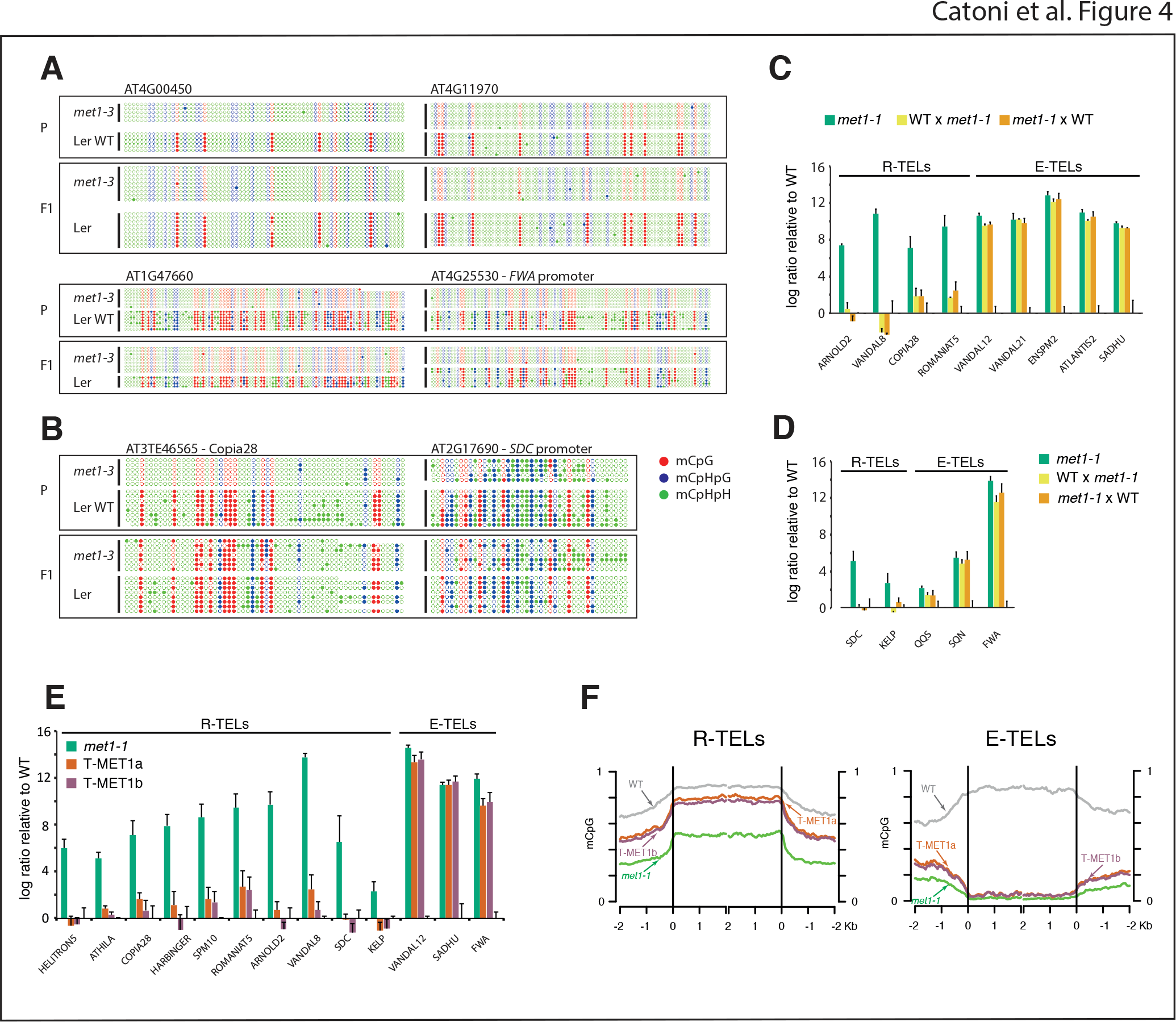
Transgenerational epiallelic stability of GELs, E-TELs and R-TELs in backcrosses.

(A) DNA methylation analyses at two GELs (ATG4G00450 and AT4G11970) and two E-TELs (AT1G47660 and AT4G25530-*FWA* promoter) of parental strains (*met1-3* mutant, which is in the Col-0 accession and Landsberg erecta (Ler) accession (panels marked P) and F1 hybrid (panels marked F1) containing *met1-3* and Ler derived chromosomes, that can be distinguished by DNA polymorphism. Open circles represent unmethylated cytosine and closed circles methylated. The colours correspond to methylation in different sequence contexts explained in (B).

(B) DNA methylation analyses at two R-TELs (AT3TE46565-Copia28 and AT2G17690-SDC promoter). General marking as in (A)

(C) Transcript levels in *met1-1* and in reciprocal F1 hybrids of *met1-1* and wild type (colour code above the graphs) of transposons assigned to R-TEL or E-TEL. The values are log ratio relative to the wild type adjusted to 0 (baseline). Graphs represent mean values of three biological repetitions, ± s.d. marked by black lines.

(D) Transcript levels in *met1-1* and in reciprocal F1 hybrids of *met1-1* and wild type of genes assigned to R-TEL or E-TEL.

(E) Transcript levels in *met1-1* and in two independent transgenic lines with a reintroduced copy of the MET1 gene (T-MET1a and T-MET1b). Colour code is given next the graphs. The identities of the tested loci are provided below the graph. General marking as in (C).

(F) Kernel density plots of CpG methylation levels in wild type (WT), *met1-1*, and complemented *met1-1* lines (T-MET1a and T-MET1b) at R-TELs and E-TELs applied to TAIR10 annotation and aligned at the 5′ end or the 3′ ends (vertical lines). Average methylation levels for each 200-bp window are plotted. Chromosome 5 was excluded from the analyses. Corresponding plot for GELs is displayed on Figure S16.

Importantly, a backcross of *met1-1* to wild type brings hypomethylated and wild-type epialleles together in the F1, which opens the possibility of trans interactions between epialleles (Greaves et al. 2012; Rigal et al. 2016). To avoid for the first time such confounding epiallelic trans-interactions, we introduced a *MET1* transgene into *met1-1* plants and examined R-TELs and E-TELs in two complemented transgenic lines with wild type-like MET1 protein levels (Figure S15 and S16). Similar to what was observed in *met1*/*MET1* F1 hybrids and in *MET1^+^* plants, R-TELs and E-TELs displayed their previously documented transcriptional attributes in both complemented transgenic lines (Figure 4E). Finally, we examined genome wide DNA methylation profiles of the two *MET1*complemented lines (Table S1 and Figure S1) and observed that transgenic MET1-dependent DNA remethylation occurs exclusively at R-TELs, and not at E-TELs or GELs (Figure 4F and S16A, B). Likewise, epigenetic proprieties of certain TEs recorded in *MET* plants (Figure S14) are consistent with those observed in the transgenic lines (Figure S16C). Taken together, these results confirmed that distinct DNA sequence properties rather than trans-epiallelic interactions determine DNA epigenetic stability or switching.

Finally, we wanted to know whether our predictive rules of DNA sequence properties in defining susceptibility to epigenetic switching have broader applicability. For that, we identified GELs, E-TELs and R-TELs in data from published experiments. For example, it has been demonstrated that progeny of heterozygous *met1-3* plants experience CpG methylation loss in the absence of MET1 during post-meiotic divisions of the haploid gametophytes (Saze et al. 2003). Therefore, it can be predicted that inbreeding of *met1-3* heterozygous plants will result in gradual methylation losses. However, since MET1 would be present at each generation during somatic development, R-TELs would be subjected to remethylation, while GELs and E-TELs would remain hypomethylated. To test this prediction, we scored methylation levels of GELs, E-TELs and R-TELs in previously reported methylation profiles of plants propagated as heterozygous *met1-3*(*metl-3*+/-) and wild-type plants (*metl-3* +/+) segregating from these lines (Stroud et al. 2013). Consistent with our predictions, complete loss of CpG methylation at GELs and strong reduction in E-TELs were found in these datasets. In contrast, levels of methylation at R-TELs were similar to wild type (Figure 5A).

**Figure 5.**
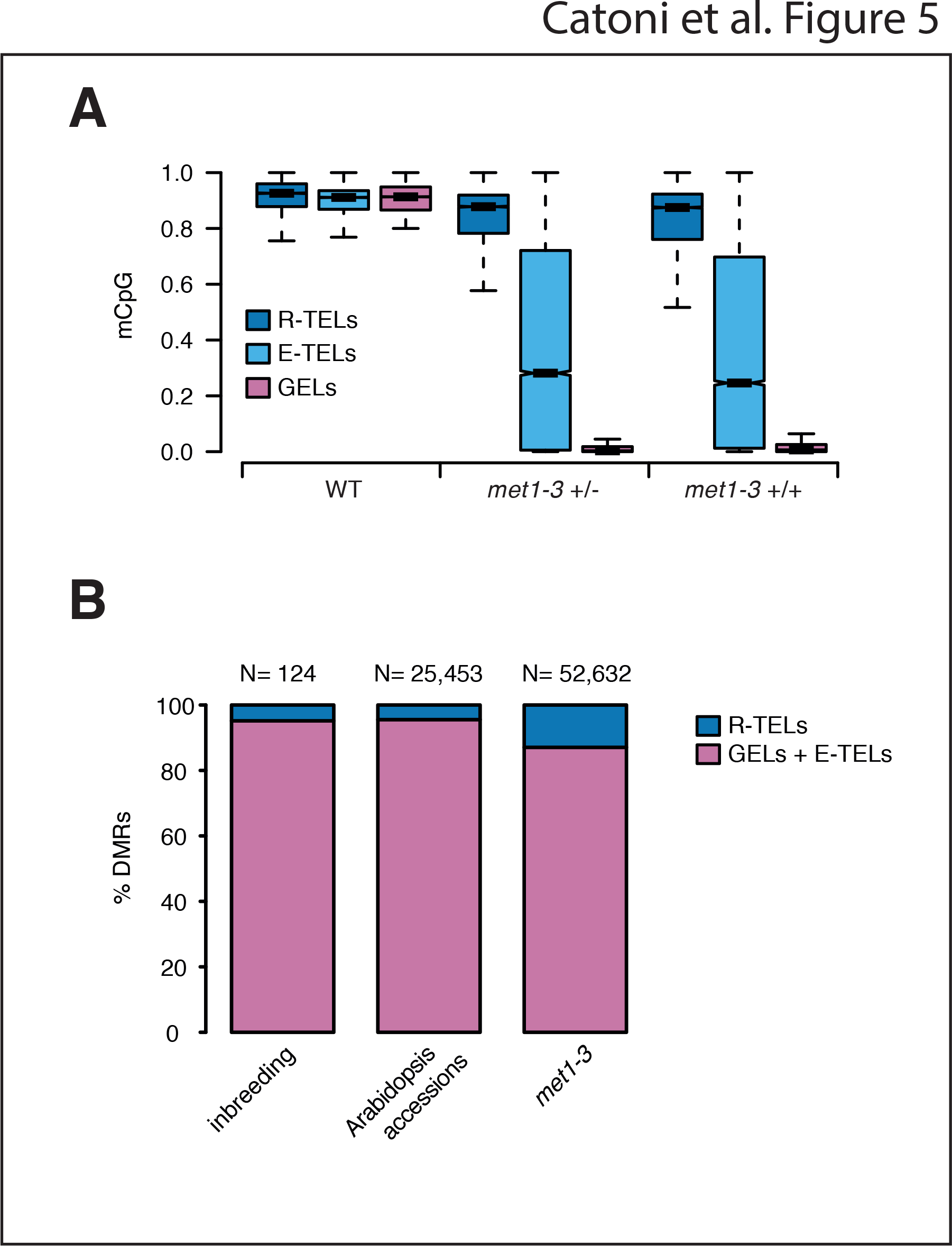
Epiallelic frequencies in Arabidopsis.

(A) Box plot of CpG methylation levels in 200-bp genomic tiles sorted as R-TELs, E-TELs and GELs in wild type, inbred *met1-3* heterozygous mutant (*met1-3* +/-) and wild type plants segregated from inbred *metl-3* heterozygous mutant (*met1-3* +/+) (Stroud et al. 2013). Only GEL tiles with on average CpG methylation ≥80% were considered for this analysis.

(B) Relative distribution of CpG DMRs (among sequences with ≥60 % CpG methylation in wild type), found in *metl-3* in long time inbreeding (Becker et al. 2011) and among Arabidopsis accessions (Schmitz et al. 2013), divided to R-TELs or GELs/E-TELs. Total number of considered DMRs (*n*) for each experimental system is given above the graphs.

Next, we tested distribution of DMRs in R-TELs, GELs and E-TELs in the results of inbreeding experiments of Arabidopsis for 30 generations (Becker et al. 2011) and in natural Arabidopsis accessions (Schmitz et al. 2013). In both populations, we registered the occurrence of DMRs at GELs and E-TELs at a high frequency and at R-TELs at a very low frequency (Figure 5B). Thus, the revealed DNA sequence properties can be used to predict transgenerational epigenetic inheritance in nature.

To test whether these DNA sequence rules are universal and thus applicable to other plant species, we turned to methylation data from recently characterized *met1* mutants of rice (Hu et al. 2014). In this mutant, CpG methylation is reduced by 76%, closely resembling levels in Arabidopsis *metl-l*(Figure S17A and Table S1). CpG methylation at rice GELs was lost efficiently and TELs were characterized by heterogeneous methylation losses (Figure 6A), which mirrored Arabidopsis results (Figure 2A, S4). We then compared the DNA sequences of Arabidopsis and rice E-TELs and R-TELs (Figure 3C, S8). Since Arabidopsis and rice genomes have different sizes, CG contents and number of repeats, we normalized their sequences according to the general properties of each genome. The characteristics of DNA sequences (specific CpG frequencies) of rice E-TELs and R-TELs and degrees of sequence repetitiveness were found to be similar to those of Arabidopsis (Figure 6B and C and S17B and C).

**Figure 6.**
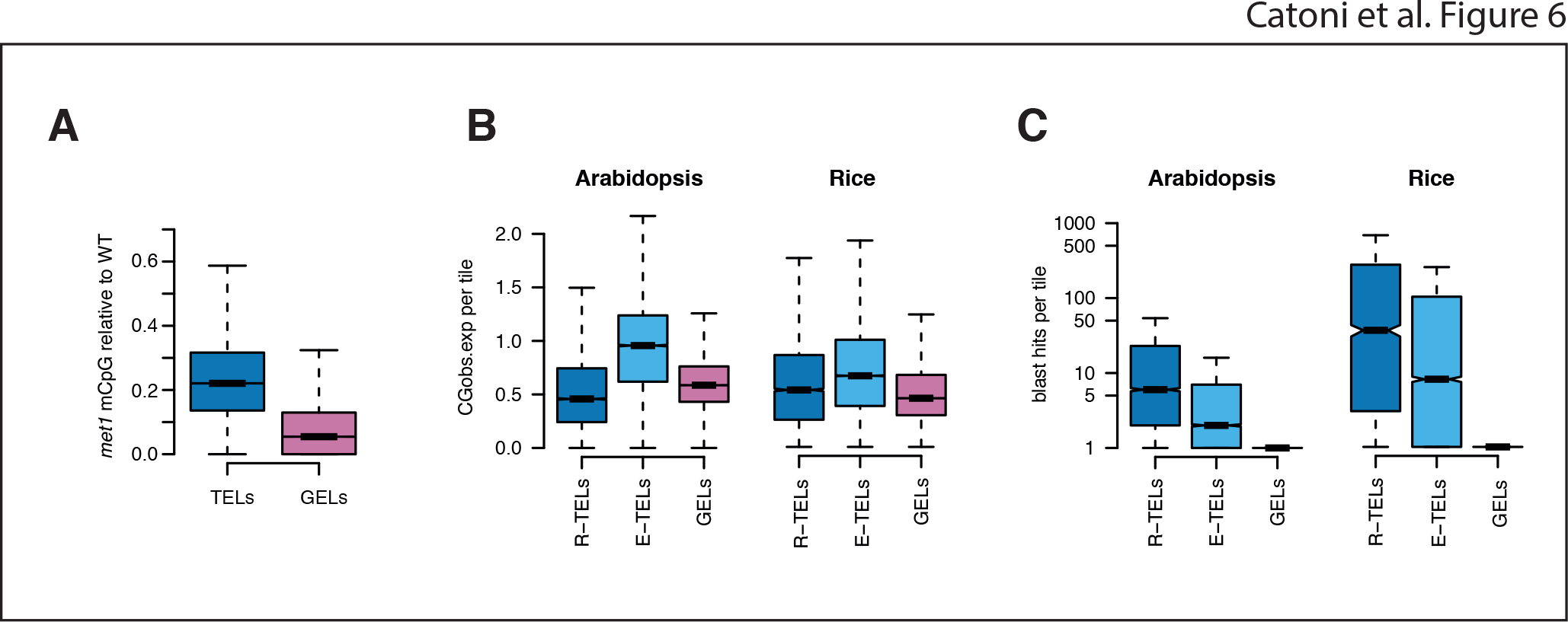
GELs, E-TELs and R-TELs in rice.

(A) Box plots of CpG methylation levels in 200-bp genomic tiles sorted as TELs and GELs of rice *MET1* relative to wild type.

(B) Box plots representing the copy number (blast hits) of 200-bp tiles assigned to R-TELs, E-TELs and GELs, comparison of Arabidopsis and rice. Each assigned tile was blasted against the corresponding reference genome and number of blast hits for each category (Blastn expectation value <10) is reported.

(C) Box plots representing the relative proportions of numbers of CpGs for R-TELs, E-TELs and GELs and their expected frequency in 200-bp tiles of the Arabidopsis and rice genomes.

It has often been discussed whether and, if so, to what extent DNA sequence itself impacts on epigenetic regulation (Schübeler 2015). Recent transgenic experiments, with site-directed insertion of various DNA fragments into a predefined chromosomal position of mammalian cells, provided evidence that the base composition of DNA sequence may attract or prevent DNA methylation according to CpG content and the occurrence of transcription factor binding sites (Krebs et al. 2014). However, molecular determinants that distinguish between chromosomal loci that rapidly revert to one dominant epigenetic state and loci that form alternative transgenerationally heritable epiallelic states (epialleles) remained largely unknown. Our results demonstrate that the properties of DNA sequence can be used to predict the epiallelic behaviour of certain classes of plant chromosomal loci. Repetitiveness and relative scarcity of CpGs are associated with rapid reversion to one dominant epigenetic state and low copy number and high CpG content are linked to the formation and support of the transgenerational stability of alternative epiallelic states.

## Methods

### Plant growth and material

Plants used in this work were derived from *Arabidopsis thaliana* Columbia-0 lines, 13^th^ and 2^nd^ generation homozygous *metl-l* (Kankel et al. 2003) and *met1-3* (Saze et al. 2003) have been used, respectively. The *met2,3,4* triple mutant was generated by crossing the lines SALK_010893, SALK_099592 and SALK_098878, genotyping and selecting the segregating homozygous mutants through PCR. Seeds were stratified at 4^°^C for 4 days and plants grown in 1/2 MS 1.5% agar vertical plates or in soil, depending on the analysis. Plants were grown under long-day conditions (21^°^C, 16 h light, 8 h dark).

### Genomic data

Histone methylation data and small RNA data for Col-0 and *met1-3* were GSE370775 and GSE10967, respectively (Deleris et al. 2012; Lister et al. 2008). DMRs recovered in the long-term inbreeding experiments and DMRs found in Arabidopsis accessions were retrieved from previously published work(Schmitz et al. 2013; Becker et al. 2011).

### Nucleic Acid Extraction

DNA and RNA were extracted from pools of 2-week-old seedlings (25-30 plants per pool) grown on plates of wild type, *met1-1* and *met1-3*. Due to epiallelic segregation in *MET1^+^* plants, two *MET1^+^*pools (8 plants each) were generated starting from genotyped MET1 wild-type leaves derived from the segregating progeny of two F1 plants, obtained by backcrossing *met1-3* with Col-0. For each genotype, genomic DNA was extracted using the Qiagen Plant DNeasy kit (Qiagen), starting from nuclei purified as described in (Becker et al. 2011). Total RNA was extracted using the Trizol (Invitrogen) method according to the manufacturer’s instructions.

### Library Preparation

Bisulfite-converted DNA libraries of *met1-1*,w*met1-3*, two *MET1^+^* replicates, two transgenic complemented lines (T-MET1a and T-MET1b) and wild type Col-0 control for genomic sequencing were performed starting from 0.5 - 1 µg of genomic DNA using the NEBNext DNA Sample PrepReagentSet1 (New England Biolabs), following the Illumina Genomic Sample Prep Guide (Illumina), as described in (Becker et al. 2011). Libraries for RNA expression analysis were prepared in triplicate from 3 µg of total RNA, processed with the GeneChip Whole Transcript Amplified Double-Stranded Target Assay (Affymetrix), according to the manufacturer’s protocol, to generate labelled cDNA for tiling microarray hybridization.

### Analysis of Gene Expression

The labelled cDNA was hybridised to the GeneChip Arabidopsis Tiling 1.0R array (Affymetrix) and scanned following the manufacturer’s instructions. Tiling array hybridisation data were processed with the R statistical software (r-project.org) and BioConductor (www.bioconductor.org) applying the chip definition file (CDF), kindly provided by Naouar et al (2009), as previously described (Yokthongwattana et al. 2010).

For real-time qRT-PCR analysis, total RNA (2 µg) was treated with RQ1 DNase (Promega) and reverse transcribed with the SuperScript VILO cDNA Synthesis Kit (Invitrogen) according to the manufacturer’s instructions. PCR reactions were carried out in triplicate using 10 ng of template cDNA, 200 nM target specific primers (Table S5), and LightCycler 480 SYBR Green I Master (Roche) in the LightCycler 480 II detection system (Roche) in a volume of 10 µl.

### Sequencing and processing

Bisulfite-converted libraries were sequenced with 2 × 101-bp paired-end reads on an Illumina GAIIx or Illumina NextSeq 500 instrument. For image analysis and base calling, we used the Illumina OLB software version 1.8. The raw reads were trimmed using Trimmomatic (Bolger et al. 2014) in order to remove adapter sequences. Reads with an average quality value of at least 15 in a window of 4 nucleotides were trimmed from both ends. After trimming, reads shorter than 16 bases were discarded. The remaining sequences (on average 77% of raw reads) were aligned against the *Arabidopsis thaliana* genome TAIR10 version using Bismark (Krueger and Andrews 2011). Duplicated reads were collapsed into one read. Chloroplast sequences were used to estimate the bisulfite conversion (on average above 99%) (Table S1). To account for non-converted DNA, we applied a correction according to (Lister et al. 2013). The number of methylated reads were decreased as: m^*^=max(0, m - nc) (where m^*^ is the corrected number of methylated reads, m is the raw number of methylated reads, n is the total number of reads and c is the conversion rate). DMRs (differentially methylated regions) were defined comparing methylation in wild type Col-0 with the other conditions analysed using the R package “DMRcaller” (Zabet and Tsang 2015). We used “noise filter” method to compute CpG and CpHpG DMRs. Briefly, the “noise filter” method uses a triangular kernel to smooth the total number of reads and the total number of methylated reads. Note that the “noise filter” method uses the assumptions of BSmooth package (Hansen et al. 2012), namely, that adjacent cytosines display correlated methylation. In particular, we used a window size of 172 nt, for CpG methylation, and 160 nt, for CpHpG methylation, to smooth the data and then we performed a Score test a each position to determine the positions that display a statistically significant differences in methylation levels between the two conditions (note that using the Score test leads to the same results as Fisher’s exact test, but was much faster to compute). At each position we computed the p-value and adjusted for multiple testing using the Benjamini and Hochberg’s method (Benjamini and Hochberg 1995) to control the false discovery and we discarded positions with FDR higher than 0.05. We further discarded the positions with less than 4 reads and the positions with differences in methylation levels lower than 0.4, in the case of CpG, and 0.2 in the case of CpHpG. Adjacent positions within 200 nt of each other were joined only if the resulting DMR displayed an minimal average number of reads per cytosine of 4 and statistically significant difference in methylation level (FDR lower than 0.05) which is at least 0.4 in the case of CpG and 0.2 in the case of CpHpG. For CpHpH DMRs, we used the “neighbouring” method, which performs the same algorithm as the “noise filter” method except that it does not perform data smoothing. In the case of CpHpHp DMRs, we considered only regions that display a minimal difference in the methylation level of 0.1 and a minimum size of 50 bp. Rice *met1* mutant and wild type control sequencing data were obtained from (Hu et al. 2014) and re-analysed with the parameters described for Arabidopsis.

### Sequence Methylation Analysis

Annotated DNA sequences or genomic 200-bp tiles (with 150 bp overlap) were classified according to the methylation information available in wild type Col-0 for Arabidopsis (provided in this work) and Nipponbare for rice (Hu et al. 2014). Methylation values were averaged among all cytosines in the same context in each TAIR10 annotated sequence or 200 bp tile, with a minimum coverage of five reads. A sequence was considered GEL if more that 5% methylation was present in CpG context, and less than 5% of methylation was present in each of the non-CpG contexts (CpHpG and CpHpH). Sequences were considered TEL if more that 5% of methylation was present in each cytosine context. Sequences were considered not methylated if less than 5% of methylation was present in each context (Table S2). Fisher exact test or Wilcoxon test were used according to the data distribution. Shapiro and Bartlett tests were used for normal distribution or variance similarity, respectively.

### Local DNA Methylation Analysis

Approximately 1 µg of DNA extracted using a Mini plant DNA kit (Qiagen) was used for bisulfite conversion with the EpiTect Bisulfite kit (Qiagen) according to the manufacturer’s instructions. One microliter of converted DNA was PCR-amplified with degenerate primers designed to match the tested target. The PCR products were cloned into pGEM-T Easy (Promega) and independent clones were sequenced for each condition analysed. KISMETH (Gruntman et al. 2008), a plant-specific tool for bisulfite sequencing analysis, was used to calculate the percentage of methylation and average different clones. We estimated a bisulfite conversion rate of 98%, based on the methylation level at the non-methylated *PHAVOLUTA* (At1g30490) locus (Bao et al. 2004).

### MET1 Antibody Production

The BAH_DMC domain of MET1 of 175 amino acids (695aa-869aa) was cloned into pDEST-15-GST (Glutathione S-Transferase tagged; Gateway, Invitrogen) using the primers met1_BAH1_attB1 and met1_BAH1_attB2 (Table S5); the transformed BL21 *E.coli* strain as used to produce the recombinant protein. Purified GST::BAH protein was injected into two rabbits (AGRO-BIO, La Ferte Saint Aubin, France). Although both rabbit sera produced non-specific signals with Arabidopsis protein extract, one serum (anti-MET-84/J77) yielded the signal expected for MET1 protein in wild type (172.4 kDa).

### Transgenic*met1-1*Complementation

The genomic version of *MET1* and the 2,677-bp upstream promoter sequence were independently amplified from genomic DNA using the primer combinations met1_adapF/met1_FLAGR, and Met1PR_adap_F/Met1PR_adap_R, respectively (Table S5). The MET1 gene was assembled into a *pGPTVII-bar-MCS* (multi-cloning-site) plasmid using the XhoI/XmaI sites and cloned into the barII_pUBQ10_MCS binary vector. The ubiquitin promoter was then removed and substituted with the *MET1* promoter using the HindIII/XhoI sites. The Agrobacterium tumefaciens pGV3101 strain was used to transform *A. thaliana met1-1* using the standard floral-dip method. Transgenic lines were selected *in vivo* for resistance to BASTA (dl-phosphinothricin, Duchefa). Two T2 lines homozygous for transgenic *MET1* gene (T-MET1a and T-MET1b) were used for preparing bisulfite-converted DNA libraries and sequenced as described for the other genotypes.

### Protein Extraction and Analysis

Arabidopsis 15-day-old seedlings grown on plates were used for protein analysis. Around 10 seedlings were pooled in 2-ml tubes and frozen in liquid nitrogen. The samples were homogenized in the TissueLyser II (Qiagen) and proteins extracted with 200 µl of 2X Laemmli Sample Buffer (Sigma). Samples were incubated for 3 min at 95^°^C and centrifuged; aliquots of15 µl were loaded onto Mini-PROTEAN precast polyacrylamide gels (Biorad) for electrophoresis and SDS-PAGE separation. The separated proteins were transferred to PVDF membrane and immunodetection performed using 1:2500 primary antibody (anti-MET-84/J77) dilution follow by 1:25000 dilution of anti-rabbit-HRP (Sigma). Signal intensities were measured with Odyssey Fc (Li-COR Biosciences).

## Acknowledgments

This work was supported by EVOBREED ERC grant 322621, Gatsby Fellowship AT3273/GLE, AENEAS and Max Planck Society.

## Author Contributions

Conception and design of the experiments: MC and JP. Nucleic acid library preparations: MC, MD, MLL, C Becker and C Bayon. Genome-wide sequencing: C Becker. Local DNA methylation profiles: JG and MC. Performance of all other experiments: MC. Analysis of the data: MC, NRZ and C Becker. Contribution of reagents/materials/analysis tools: JP and DW. Writen the paper: MC and JP.

